# Carotenoid and tocopherol fortification of zucchini fruits using a viral RNA vector

**DOI:** 10.1101/2021.02.18.431807

**Authors:** Fakhreddine Houhou, Teresa Cordero, Verónica Aragonés, Maricarmen Martí, Jaime Cebolla-Cornejo, Ana Pérez de Castro, Manuel Rodríguez-Concepción, Belén Picó, José-Antonio Daròs

## Abstract

Carotenoids and tocopherols are health-promoting metabolites in livestock and human diets. Some important crops have been genetically modified to increase their content. Although the usefulness of transgenic plants to alleviate nutritional deficiencies is obvious, their social acceptance has been controversial. Here, we demonstrate an alternative biotechnological strategy for carotenoid and tocopherol fortification of edible fruits in which no transgenic DNA is involved. A viral RNA vector derived from *Zucchini yellow mosaic virus* (ZYMV) was modified to express a bacterial phytoene synthase (crtB), and inoculated in zucchini (*Cucurbita pepo* L.) leaves nurturing pollinated flowers. After the viral vector moved to the developing fruit and expressed *crtB*, the rind and flesh of the fruits developed yellow-orange rather than green color. Metabolite analyses showed a substantial enrichment in health-promoting carotenoids, such as α- and β-carotene (pro-vitamin A), lutein and phytoene, in both rind and flesh. Considerably higher accumulation of α- and γ-tocopherol was also detected, particularly in fruit rind. Although this strategy is perhaps not free from controversy due to the use of genetically modified viral RNA, our work does demonstrate the possibility of metabolically fortifying edible fruits using an approach in which no transgenes are involved.

## Introduction

Plants produce a vast number of secondary metabolites, many of them of great interest in food, pharmaceutical and industrial applications. However, valuable plant metabolites are not infrequently produced in limited amounts or without the desired chemical properties. The ability to manipulate endogenous metabolic pathways or to deploy novel pathways in plants via biotechnological metabolic engineering approaches should contribute to overcoming these issues and successfully establishing these autotrophic organisms as “green factories” able to sustainably provide most of the molecules required by humankind. In contrast to other systems, production in plants is cheap, easily scalable, mainly fueled by sunlight, and, if properly managed, free of human pathogens.

In fact, plants are currently being used as platforms to produce recombinant proteins and peptides in both stable genetic transformation or transient expression approaches ^1,2^. Stable genetic transformation of plants is a labor-intensive and time-consuming process that frequently leads to results with undesired variability. By contrast, transient expression systems, such as those that employ *Agrobacterium tumefaciens*, viral vectors, or a combination of both, offer a rapid alternative for reaching some biotechnological goals. We have recently showed that plant virus-derived vectors can be used not only to produce recombinant proteins, but also to engineer plant metabolism in cases in which the expressed proteins are regulatory factors or biosynthetic enzymes that interact with the natural host plant metabolism. Virus-based expression of transcription factors Delila and, particularly, Rosea1 from *Antirrhinum majus* has been shown to lead to massive accumulation of anthocyanins in plant tissues ^3,4^. Similarly, virus-based expression of *Pantoea ananatis* phytoene synthase (crtB), the enzyme catalyzing the first step of carotenoid biosynthesis in this soil bacterium, has been demonstrated to lead to a substantial accumulation of phytoene and other carotenoids in plant tissues ^5,6^. Virus-based co-expression of crtB and a *Crocus sativus* carotenoid cleavage dioxygenase (CCD2L) induced large accumulation in *N. benthamiana* leaves of the apocarotenoid crocins and picrocrocin, which naturally accumulate in saffron stigma and are main constituents of the valued spice ^7^. However, these achievements were mainly produced in model plants, such as *Nicotiana tabacum*, *N. benthamiana* or *Arabidopsis thaliana*. Our goal here was to translate this ability to produce health-promoting metabolites such as carotenoids to edible tissues, particularly edible fruits. Generally, animals do not biosynthesize carotenoids, which are essential nutrients that must be acquired from diet ^8^. Consequently, highly relevant projects aimed to the carotenoid fortification of important crops have been undertaken in last two decades ^9–12^. To this end, we investigated a strategy to induce accumulation of carotenoids in edible zucchini (*Cucurbita pepo* L.) fruits using an RNA-based viral vector derived from *Zucchini yellow mosaic virus* (ZYMV; genus *Potyvirus*, family *Potyviridae*) that expresses the bacterial biosynthetic enzyme crtB.

## Results and Discussion

### Inoculation of zucchini plants with ZYMV-crtB at the right developmental stage produces induces yellow-orange fruits

We previously observed that inoculation of zucchini plants with a ZYMV-based vector that expressed *P. ananatis* crtB (ZYMV-crtB) caused infected leaves to turn bright yellow ^5^. We have, more recently, understood that this phenotype is a consequence of the heterologous crtB enzyme inducing phytoene accumulation beyond a threshold that triggers transformation of leaf chloroplasts into chromoplasts, which is accompanied by the accumulation of high levels of downstream carotenoids ^6^. To test whether we could trigger the chloroplast-to-chromoplast transformation in fruits, with concomitant carotenoid overaccumulation, we grew seedlings of a zucchini cultivar (MU-CU-16) ^13,14^, which produces marketable dark green uniformly cylindrical fruits, and mechanically inoculated the plants (n=3) with ZYMV-crtB at one-week intervals (Fig. 1A). These intervals corresponded to five different developmental stages (Fig. 1B): plants with (1^st^) male buds, (2^nd^) female buds, (3^rd^) male flowers at anthesis, (4^th^) female flowers at anthesis, and (5^th^) female flowers one week after the anthesis, with fruits of approximately 20 cm.

**Fig. 1.**
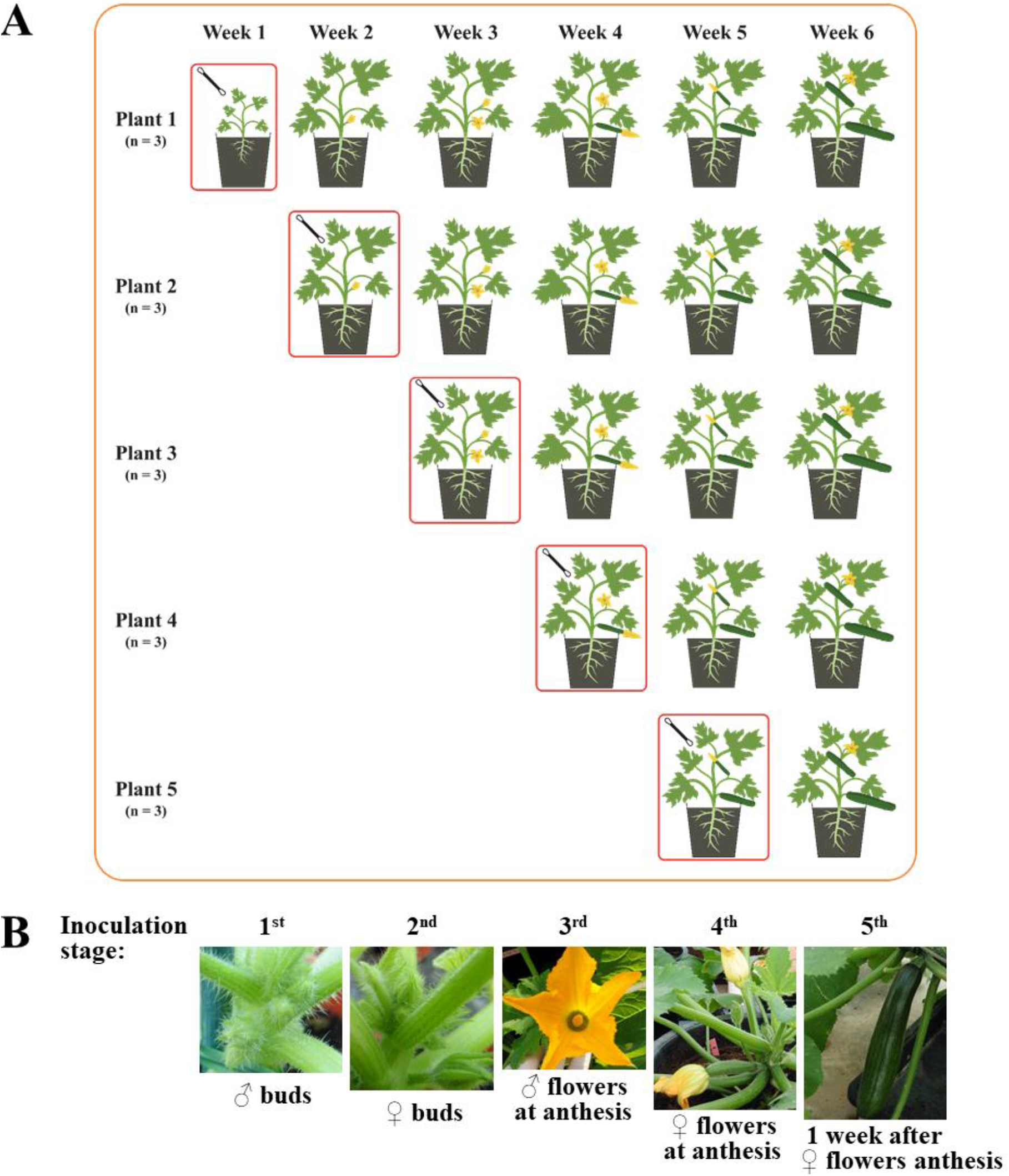
Zucchini plant inoculation process with viral vector ZYMV-crtB. (A) Plants, in groups of three, were mechanically inoculated (red rounded rectangle) at one-week intervals, such that each group became infected at a different developmental stage. (B) Pictures of flower and fruit developmental stages at which plants were inoculated.

All inoculated plants showed the first symptoms of infection at approximately 7 days post-inoculation (dpi) and symptomatic tissue turned yellow during the following days (Fig. 2A), as previously reported ^5^. Interestingly, the development of flowers and fruits of infected plants was very different depending of plant developmental stage at inoculation time. Plants inoculated at the male bud stage (1^st^) did not develop female flowers. In contrast, those inoculated at the female bud stage (2^nd^) resulted in completely yellow female buds (Fig. 2B), although flowers stopped growing before reaching anthesis. Zucchini plants inoculated at the male flower anthesis (3^rd^ stage) completed flower development and female flowers reached anthesis (Fig. 2C). However, fruits did not continue growing after pollination (Fig. 2D). Despite immature, these plants produced different types of fruits. Those below the inoculated leaves showed less yellow pigmentation than those above the inoculated leaf (Fig. 2E). Interestingly, plants inoculated having female flowers at anthesis (4^th^ stage) developed fruits after pollination, which displayed different degrees of green to yellow-orange color in their rinds (Fig. 2F), having a uniformly orange flesh (Fig. 2G). Plants inoculated one week after female flower anthesis (5^th^ stage) developed fruits of commercial size. These fruits remain with dark green rinds or developed orange speckles (Fig. 2H), but remarkably they exhibited a uniformly orange flesh (Fig. 2I). Taken together, these results suggest that a viral RNA vector can be used to specifically express a phytoene synthase, such as the bacterial crtB, in the fruits of adult zucchini plants to transform their color from green to yellow-orange. Importantly, although pigment accumulation is achieved at different developmental stages, plants with female flowers at or after the anthesis must be inoculated to produce fruit of commercial size with orange flesh.

**Fig. 2.**
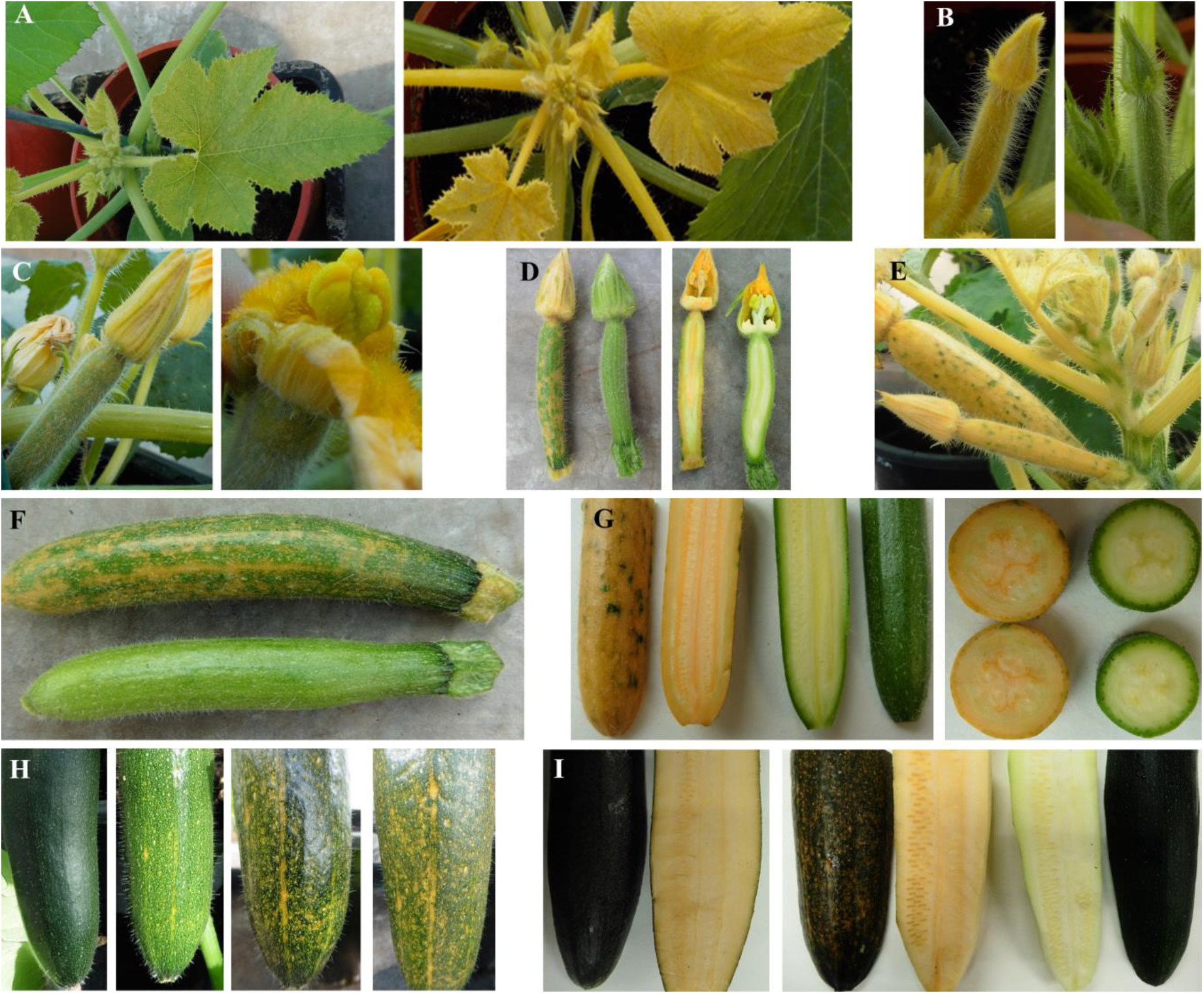
Photographs of zucchini plants inoculated with ZYMV-crtB. (A) Plants inoculated at 1^st^ stage (male buds) that exhibit the first symptoms of infection and a bright yellow pigmentation at 7 and 14 dpi, respectively. (B) Yellow and green female buds from plants inoculated at 2^nd^ stage (female buds) and a non-inoculated control, respectively. (C) Female flowers with yellow speckles at preanthesis (left) and anthesis (right) from plants inoculated at 3^rd^ stage (male flowers at anthesis). (D) Preanthesis female flower from a plant inoculated at 3^rd^ stage (left), compared to a non-inoculated control (right). Longitudinal sections of the same flowers are also shown. (E) Female flowers from a plant inoculated at 3^rd^ stage, developed over the inoculated leaf, exhibiting intense yellow pigmentation. (F) Fruit (upper) from a plant inoculated at 4^th^ stage (female flowers at anthesis), compared to a non-inoculated control (lower) at 14 dpi. (G) Details of the uniformly orange flesh of a fruit from an inoculated plant (left) compared to a control (right), at 14 dpi. (H) Fruits of commercial size from plants inoculated at 5^th^ stage (one week after anthesis of female flowers). From left to right, non-inoculated control and fruits exhibiting different degrees of yellow-orange speckles at 5, 10 and 15 dpi. (I) Longitudinal sections of fruits from inoculated plants (left) that exhibited orange flesh compared to a control (right).

### Pigmented fruits from inoculated plants accumulate the viral vector

If pigmentation of zucchini fruits from inoculated plants results from viral vector-based expression of the bacterial crtB, these fruits, in contrast to green zucchinis from non-inoculated controls, must contain the recombinant virus ZYMV-crtB. To confirm this prediction, we purified RNA from fruit samples of three predominantly yellow and three speckled zucchinis, each of them harvested from a different inoculated plant. As a control, we also purified RNA from three green zucchinis from three independent non-inoculated plants. Reverse transcription (RT)-polymerase chain reaction (PCR) amplification of a cDNA corresponding to ZYMV coat protein cistron using specific primers demonstrated the presence of the virus in both the speckled and yellow zucchinis, but not in the green non-inoculated controls (Fig. 3).

**Fig. 3.**
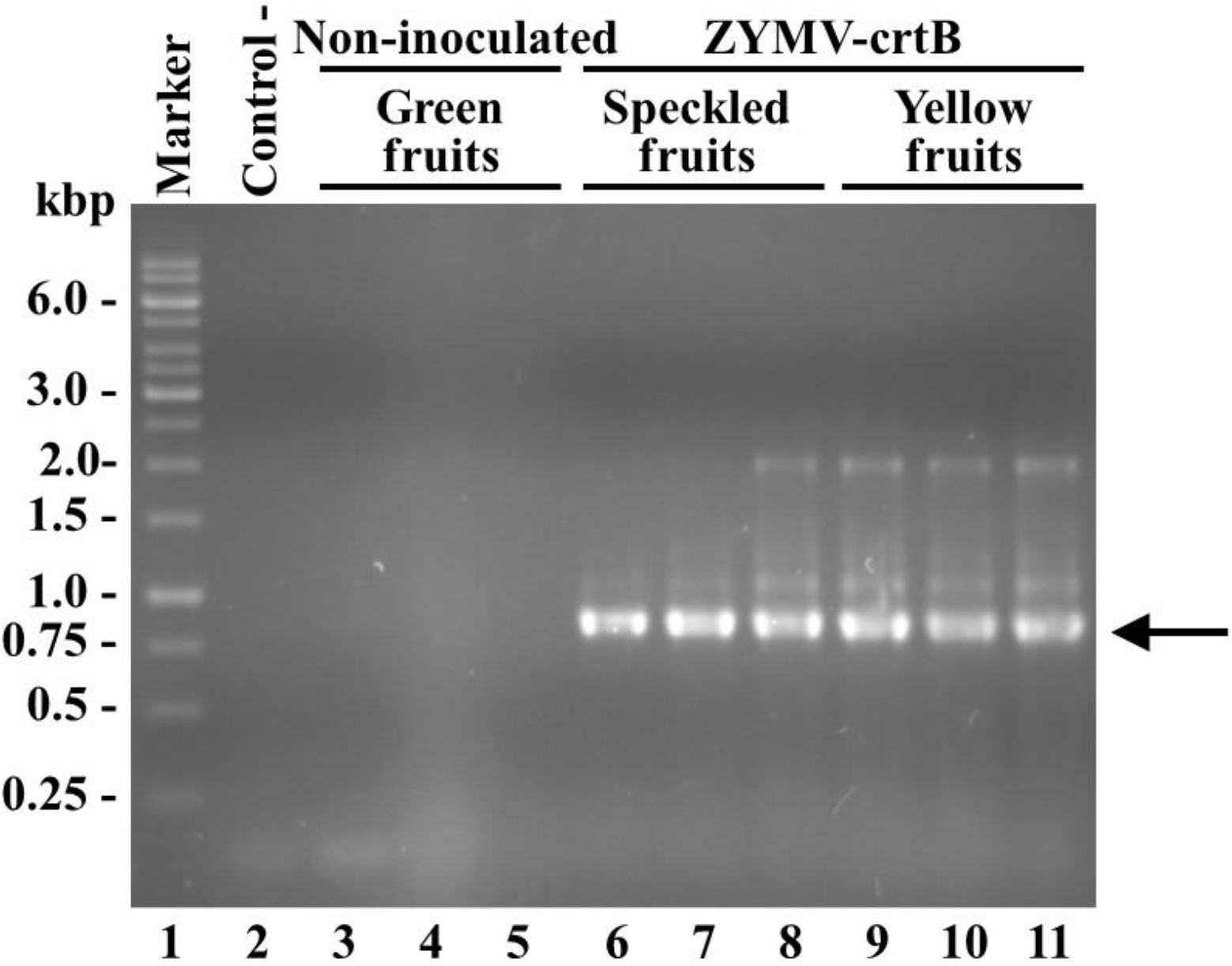
Diagnosis of ZYMV-crtB in zucchini fruits. RNA was purified and the ZYMV CP cistron amplified by RT-PCR. Products were separated by electrophoresis in a 1% agarose gel that was stained with ethidium bromide. Lane 1, DNA marker with some sizes (in kbp) on the left; lane 2, RT-PCR control with no RNA added; lanes 3 to 5, amplification products from green fruits harvested from non-inoculated plants; lanes 6 to 8 and 9 to 11, amplification products from speckled (lanes 6 to 8) and fully yellow fruits (lanes 9 to 11) harvested from ZYMV-crtB-inoculated plants. Band corresponding to ZYMV CP cistron is indicated by an arrow on the right.

### Pigmented fruits from inoculated plants exhibit increased carotenoid accumulation

Another prediction was that the pigmented fruits produced by inoculation with ZYMV-crtB exhibited an increase in carotenoid accumulation. To confirm this, we extracted carotenoids from green and pigmented fruits harvested, respectively, from non-inoculated and ZYMV-crtB-inoculated plants and quantified them after separation by high performance liquid chromatography (HPLC). Analysis was performed at the fruit commercial ripening stage, but also at two earlier developmental stages: fruit at post-anthesis (medium size) and fruit at pre-anthesis (small size), sampled respectively from plants inoculated at 5^th^, 4^th^ and 3^rd^ stages. Four representative carotenoids were quantified, namely phytoene, α- and β-carotene, and lutein. Remarkably, compared to the green controls from non-inoculated plants, pigmented fruits from ZYMV-crtB-inoculated plants, at all three developmental stages, exhibited a dramatically increased accumulation of the four analyzed carotenoids in whole fruits, as well as in rind and flesh tissues (Fig. 4 and Table 1). Carotenoids provide some of the most important nutritional benefits of consuming squash. More specifically, β-carotene and lutein are the most abundant carotenoids in the fruits of *C. pepo*, although these carotenoids are only abundant in those fruits with yellow-orange flesh that are consumed when fully mature ^15,16^. Zucchini fruits are consumed immature and naturally contain low amounts of these two carotenoids.

**Fig. 4.**
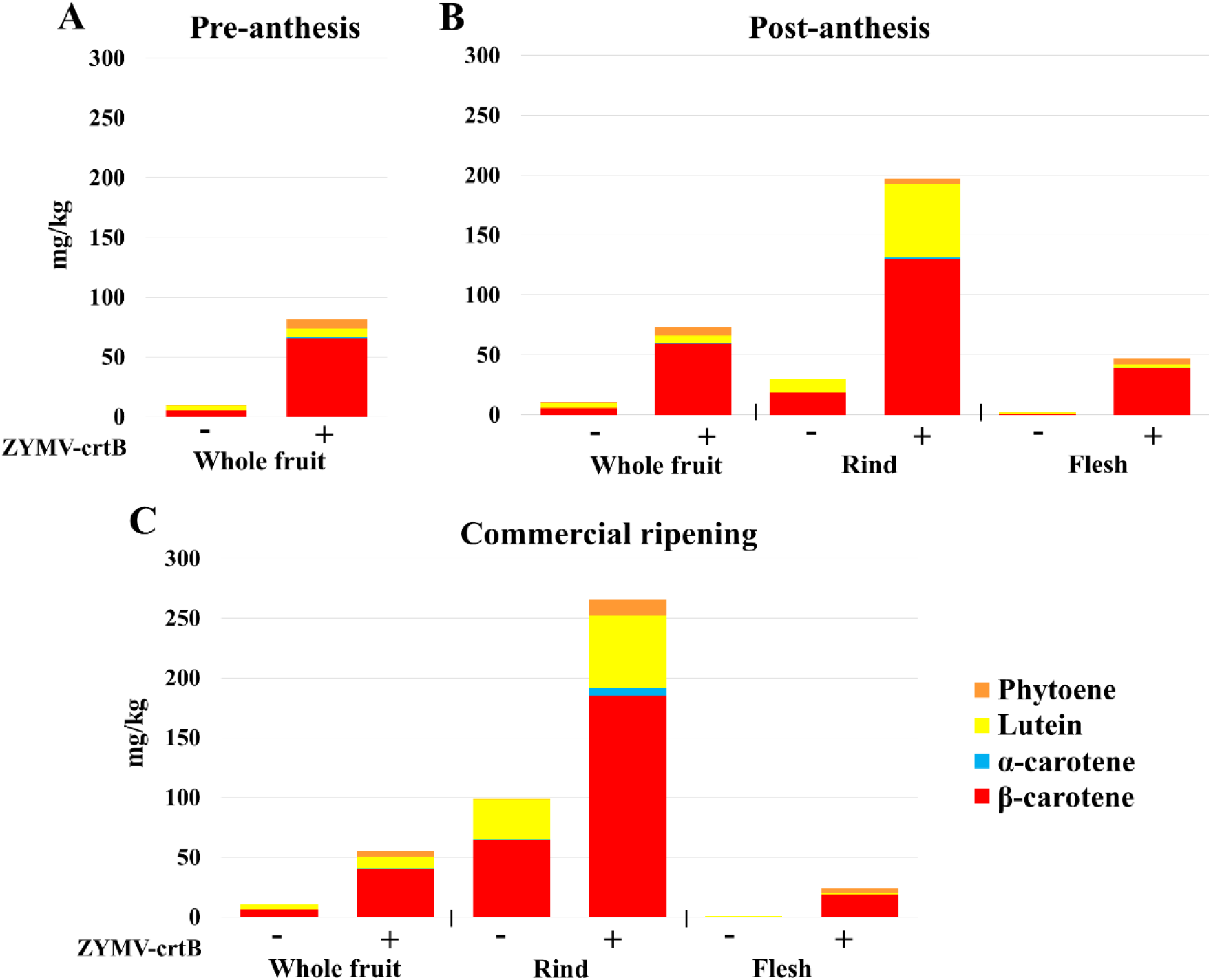
Accumulation of some representative carotenoids in whole fruit, rind, and flesh (as indicated) of green and yellow-orange zucchini fruits harvested from non-inoculated (-) and ZYMV-crtB-inoculated (+) plants, respectively, at three developmental stages. (A) Pre-anthesis, (B) post-anthesis, and (C) commercial ripening. Bars represent average accumulation in three fruits harvested from three different plants. Numerical quantifications with standard deviations are in Table 1.

**Table 1.** Carotenoid content (mg·kg^−1^ fresh weight; mean ± standard deviation, n = 3) of zucchini fruits at different stages of development in non-inoculated controls and ZYMV-crtB-inoculated plants. For the commercial ripening and post-anthesis fruit sizes, peel and pulp were also separately analyzed.

| Fruit size | Pre-anthesis (small size) |  |  |  |  |  |
| --- | --- | --- | --- | --- | --- | --- |
| Tissue | Whole fruit |  |  |  |  |  |
| Treatment | Control | Inoculated |  |  |  |  |
| β-carotene | 5.41±0.71 | 65.75±2.52 |  |  |  |  |
| α-carotene | 0.00±0.00 | 0.99±0.08 |  |  |  |  |
| Lutein | 4.27±0.59 | 7.03±0.32 |  |  |  |  |
| Phytoene | 0.02±0.03 | 7.73±0.34 |  |  |  |  |
| Fruit size | Post-anthesis (medium size) |  |  |  |  |  |
| Tissue | Whole fruit |  | Rind |  | Flesh |  |
| Treatment | Control | Inoculated | Control | Inoculated | Control | Inoculated |
| β-carotene | 5.67±0.61 | 58.97±3.60 | 18.23±1.98 | 129.63±0.50 | 0.13±0.08 | 38.67±6.15 |
| α-carotene | 0.00±0.00 | 0.82±0.11 | 0.00±0.00 | 1.83±0.02 | 0.00±0.00 | 0.32±0.08 |
| Lutein | 4.30±0.60 | 6.26±1.28 | 11.82±0.93 | 60.64±0.20 | 1.44±0.20 | 2.69±0.39 |
| Phytoene | 0.07±0.01 | 7.08±0.56 | 0.00±0.00 | 5.05±0.00 | 0.15±0.04 | 5.29±1.53 |
| Fruit size | Commercial ripening |  |  |  |  |  |
| Tissue | Whole fruit |  | Rind |  | Flesh |  |
| Treatment | Control | Inoculated | Control | Inoculated | Control | Inoculated |
| β-carotene | 6.35±2.11 | 39.78±5.17 | 64.54±34.92 | 184.93±59.03 | 0.05±0.09 | 18.83±1.35 |
| α-carotene | 0.00±0.00 | 1.26±0.15 | 0.32±0.37 | 6.89±2.59 | 0.00±0.00 | 0.21±0.03 |
| Lutein | 4.54±1.00 | 9.51±1.41 | 33.56±16.94 | 60.35±31.04 | 0.99±0.08 | 1.22±0.05 |
| Phytoene | 0.00±0.00 | 4.44±0.15 | 0.04±0.05 | 13.37±4.14 | 0.00±0.00 | 3.71±0.96 |

Fruits from ZYMV-crtB-inoculated plants exhibited up to 12-fold increase in β-carotene content relative to control fruits. At the commercial ripening stage, pigmented whole fruits contained 6-fold higher levels of the provitamin A carotenoid compared to controls. Although β-carotene amounts were higher in rinds (peel), increases were more important in flesh (pulp), as that of control fruits accumulate insignificant amounts of this pigment. Besides being the main precursor of vitamin A, β-carotene displays antioxidant activity that inhibits DNA damage and enhances immune system ^17^. Conversely, lutein amounts were nearly doubled in whole fruits from inoculated plants at commercial ripening stage, mostly caused by a similar increase in the edible fruit rind (Fig. 4 and Table 1). Lutein is a non-provitamin A carotenoid involved in the macular pigment of the human retina and intake decreases the risk of macular degeneration and other ocular diseases ^18^. In addition to these two main carotenoids, accumulation of phytoene and α-carotene in fruits of inoculated plants was particularly relevant, since content of these two carotenoids in control fruits was negligible (Fig. 4 and Table 1). Phytoene is a health-promoting carotenoid normally absent from green fruits and vegetables, and α-carotene has provitamin A activity (although only half that of β-carotene) ^17^. Taken together, these results indicate that zucchini fruits from ZYMV-crtB-inoculated plants not only develop an attractive yellow-orange color, but are highly enriched in healthy carotenoids.

### Pigmented fruits from inoculated plants exhibit increased tocopherol accumulation

Carotenoids are synthesized from geranylgeranyl diphosphate (GGPP), a precursor that is also used for the production of phytol for chlorophylls, but also for other nutritionally relevant plastidial isoprenoids such as tocopherols (vitamin E) ^17,19,20^. The crtB enzyme is known to convert GGPP into phytoene, the first committed step of the carotenoid pathway. To test whether increased diversion of GGPP towards the carotenoid pathway might negatively impact tocopherol accumulation, the levels of these isoprenoids were also determined in the rind and whole fruit at the commercial ripening stage. ZYMV-crtB-inoculated plants exhibited not a drop but a clear increase in α- and γ-tocopherol contents, especially in the rind of the fruits (Fig. 5 and Table 2). By contrast, chlorophylls levels were slightly decreased in pigmented fruits. Chlorophyll degradation is normally associated to viral infection and it has also been observed upon virus-based expression of crtB in tobacco leaves ^5^. It is therefore possible that the phytol released upon chlorophyll breakage might be recycled to produce tocopherols in the zucchini fruits of virus-inoculated plants. As a result, vitamin E levels are enriched 5-fold in whole fruits and 6-fold in the rind. Some zucchini yellow cultivars have been obtained introgressing the *B* gene, which affects plastogenesis, bypassing the proplastid to chloroplast transition and preventing the accumulation of chlorophylls. However, these cultivars exhibit a compromised tocopherol accumulation ^21^.

**Fig. 5.**
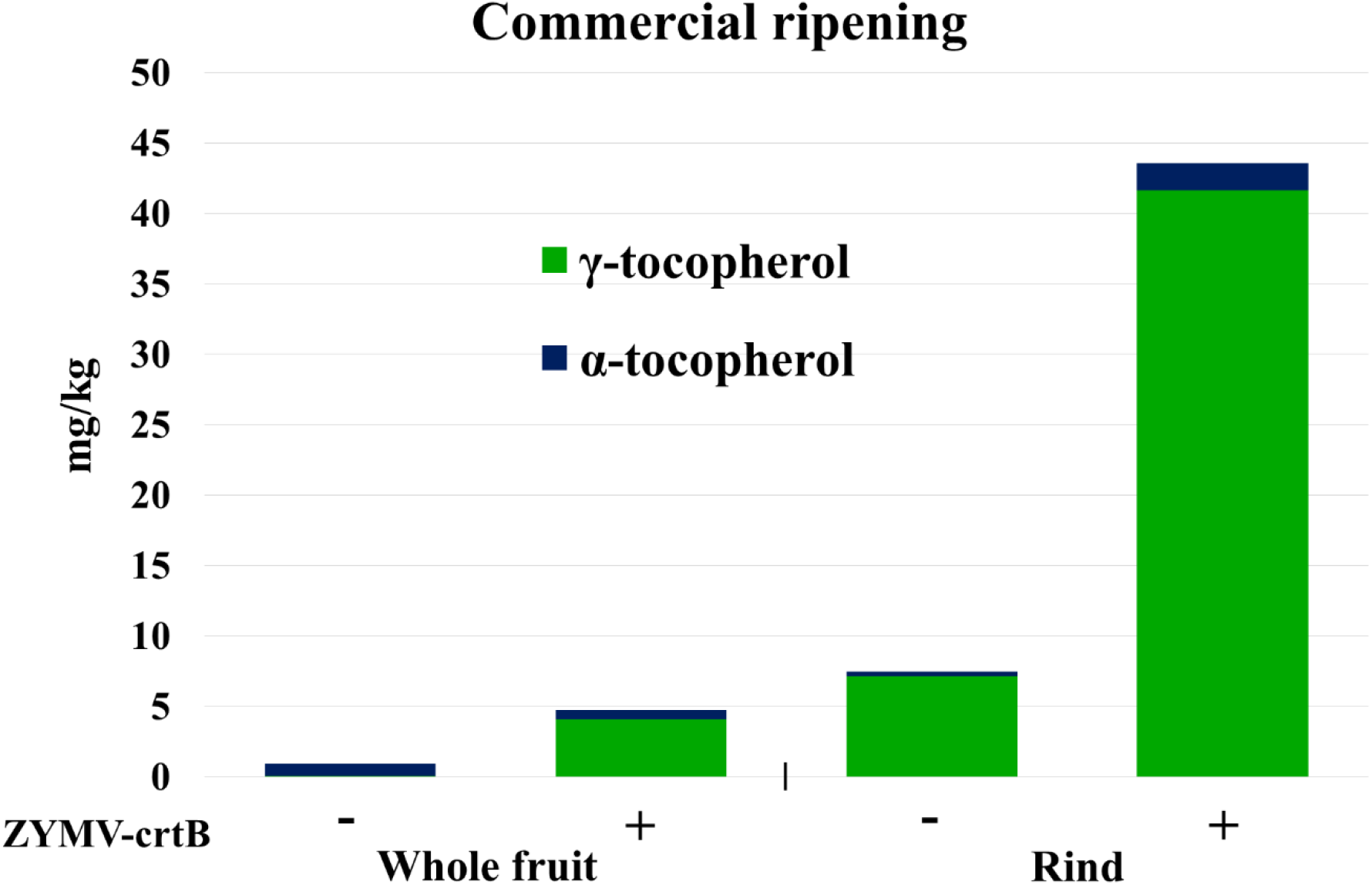
Accumulation of α- and γ-tocopherol in whole fruit and rind (as indicated) of green and yellow-orange zucchini fruits harvested from non-inoculated (-) and ZYMV-crtB-inoculated (+) plants, respectively, at commercial ripening stage. Bars represent average accumulation in three fruits harvested from three different plants. Numerical quantifications with standard deviations are in Table 2.

**Table 2.** Tocopherol content (mg·kg^−1^ fresh weight; mean ± standard deviation, n = 3) of zucchini fruits at commercial ripening development in non-inoculated controls and ZYMV-crtB-inoculated plants. Rind was also separately analyzed.

| Fruit size | Commercial ripening |  |  |  |
| --- | --- | --- | --- | --- |
| Tissue | Whole fruit |  | Rind |  |
| Treatment | Control | Inoculated | Control | Inoculated |
| γ-tocopherol | 0.04±0.07 | 4.06±0.08 | 7.13±4.92 | 41.63±5.32 |
| α-tocopherol | 0.89±0.14 | 0.67±0.48 | 0.31±0.44 | 1.92±0.42 |

Taken together, these analyses demonstrate that yellow-orange zucchinis harvested from ZYMV-crtB-inoculated plants provide nutritional benefits compared to those from market cultivars, including a substantial accumulation of highly valuable dietary isoprenoids, such as the carotenoids phytoene, lutein and others with provitamin A activity (β-carotene and α-carotene), as well as the vitamin E α- and γ-tocopherol. Also, these results show that the use of genetically modified RNA viruses as vectors is a viable strategy for metabolically fortifying the edible tissues of non-transgenic plants. As long as biosecurity issues related to the use of manipulated viruses are properly managed ^22^, this novel process presents strong potential to bring new metabolically-fortified products to the market.

### A viral RNA vector can be used to biofortify edible fruit with no genetic transformation

In conclusion, in this work we demonstrate that metabolically fortified zucchini fruits of a visually attractive yellow-orange color can be produced by using a viral RNA vector to trigger carotenoid and tocopherol overaccumulation in fruits, with no manipulation of the plant genome. This strategy is easier and faster than both classic breeding and conventional genetic engineering. The use of engineered RNA viruses in metabolic fortification of edible plant organs, subject to appropriate regulation, may contribute to bringing new products to the food, animal feed, and pharma industries.

## Materials and Methods

### Virus inoculum

One-month old zucchini plants (MU-CU-16, COMAV accession BGV004370) were infiltrated ^23^ with *A. tumefaciens* C58C1 co-transformed with the helper plasmid pCLEAN-S48 ^24^ and pGZYMV-crtB ^5^, which contains an infectious ZYMV cDNA (GenBank accession number KX499498) in which the cDNA of *P. ananatis* crtB ^25^ was inserted between the NIb and CP cistrons, flanked by sequences that complement the viral NIaPro cleavage sites ^5^. In pGZYMV-crtB, the cDNA of the recombinant virus is under control of *Cauliflower mosaic virus* (CaMV) 35S promoter and terminator. Plants were kept in a greenhouse at 25°C with a 16/8 h day/night cycle. Ten dpi, symptomatic tissue was harvested, cut into pieces approximately 50 mg, and stored at −80°C. For mechanical inoculation of zucchini plants, one of tissue piece (50 mg) was ground using a 4-mm steel ball in a 2-ml Eppendorf tube in a bead beater (VWR) for 1 min at 30 revolutions·s^−1^. Powder was brought to the bottom of the tube by a brief centrifugation, after which 1 ml of inoculation buffer (50 mM potassium phosphate, pH 8.0, 1% polyvinylpyrrolidone-10, 1% polyethylene glycol-6000, 10 mM 2-mercaptoethanol) was added and the mixture thoroughly vortexed.

### Plant inoculation

A10 μl drop of 10% carborundum in inoculation buffer was deposited on the abaxial side of a fully expanded leaf of a zucchini plant. A cotton swab soaked in virus inoculum was smoothly rubbed on the leaf surface. Plants were kept in a greenhouse at 25°C with a 16/8 h day/night cycle.

### Virus diagnosis

RNA was purified using silica gel columns (Zymo Research) from pieces of zucchini fruits. Aliquots of the RNA preparations were subjected to RT with primer PI (5’-AGGCTTGCAAACGGAGTCTAA-3’) and RevertAid reverse transcriptase (Thermo Scientific). RT products were amplified by PCR with *Thermus thermophilus* DNA polymerase (Biotools) using primers PII (5’-TCAGGCACTCAGCCAACTGTGG-3’) and PIII (5’-CTGCATTGTATTCACACCTAGT-3’). PCR products were separated by electrophoresis in a 1% agarose gel in TAE buffer (40 mM Tris, 20 mM sodium acetate, 1 mM EDTA, pH 7.2), and the gel was stained in a solution of 0.5 μg/ml ethidium bromide.

### Carotenoid and tocopherol analyses

Fruit pieces (0.3 g) were homogenized and subjected to extraction with 10 ml hexane:acetone:methanol (50:25:25) for 30 min at 4°C with continuous gentle shaking. Then, 1 ml of water was added and samples were centrifuged for 5 min. The organic layer was recovered and dried in a speedvac. The solid residue was then resuspended in ethyl acetate:acetone:methanol (20:60:20), filtered, and frozen (−80°C) until analysis. Carotenoids were quantified in an Agilent 1200 series HPLC system (Agilent Technologies, Waldbronn) using a Kinetex-XB C18 fused core column (150 mm length × 4.6 mm internal diameter, and 2.6 μm particle size) from Phenomenex (Torrance) following a published procedure ^26^, with minor modifications. Briefly, 10 μl aliquots of the samples were injected and separated using a mobile phase with two components, solvent A (acetonitrile:methanol:water, 84:9:7 v:v:v) and solvent B (methanol:ethyl acetate, 68:32 v:v). The sample was eluted using a linear gradient from 100% solvent A to 100% solvent B for 12 min; an isocratic elution of 100% B was then maintained for 7 min. Afterwards, a linear gradient to 100% solvent A was applied for 1 min, followed by an isocratic elution for 4 min to allow the column to re-equilibrate. Each sample was analyzed in duplicate. α-carotene, β-carotene, and lutein absorbance were measured at 445 nm, and phytoene absorbance at 295 nm. Tocopherols were determined in the same extract obtained for carotenoid analysis and using the same separation methodology, but in this case, detection was performed using a fluorimetric detector that worked at an excitation λ = 290 nm and emission λ = 330 nm ^27^.

## Acknowledgements

This research was supported by grants BIO2017-83184-R, BIO2017-84041-P, BIO2017-90877-REDT and AGL2017-85563-C2-1-R from Ministerio de Ciencia e Innovación (Spain), through Agencia Estatal de Investigación (cofinanced European Regional Development Fund) M.M. is the recipient of a predoctoral fellowship (FPU16/05294) from Ministerio de Educación, Cultura y Deporte (Spain). We also thank the genebank from Instituto Universitario de Conservación y Mejora de la Agrodiversidad Valenciana (COMAV), Universistat Politècnica de València.

## Author contributions

MRC, BP and JAD designed the experiments with input from the rest of the authors. FH, TC, VA, MM, JCC, APC and BP performed the experiments. All authors analyzed the results. MRC, BP and JAD wrote the manuscript with input from the rest of the authors.

## Conflict of Interest

The authors declare that they have no conflict of interest.

## Notes

### Competing Interest Statement

The authors have declared no competing interest.

